# Mutation landscape of SARS-CoV-2 reveals five mutually exclusive clusters of leading and trailing single nucleotide substitutions

**DOI:** 10.1101/2020.05.07.082768

**Authors:** Akhilesh Mishra, Ashutosh Kumar Pandey, Parul Gupta, Prashant Pradhan, Sonam Dhamija, James Gomes, Bishwajit Kundu, Perumal Vivekanandan, Manoj B. Menon

## Abstract

The COVID-19 pandemic has spread across the globe at an alarming rate. However, unlike any of the previous global outbreaks the availability of a large number of SARS-CoV-2 sequences provides us with a unique opportunity to understand viral evolution in real time. We analysed 1448 full-length (>29000 nt) sequences available and identified 40 single-nucleotide substitutions occurring in >1% of the genomes. Majority of the substitutions were C to T or G to A. We identify C/Gs with an upstream TTT trinucleotide motif as hotspots for mutations in the SARS-CoV-2 genome. Interestingly, three of the 40 substitutions occur within highly conserved secondary structures in the 5’ and 3’ regions of the genomic RNA that are critical for the virus life cycle. Furthermore, clustering analysis revealed unique geographical distribution of SARS-CoV-2 variants defined by their mutation profile. Of note, we observed several co-occurring mutations that almost never occur individually. We define five mutually exclusive lineages (A1, B1, C1, D1 and E1) of SARS-CoV-2 which account for about three quarters of the genomes analysed. We identify lineage-defining leading mutations in the SARS-CoV-2 genome which precede the occurrence of sub-lineage defining trailing mutations. The identification of mutually exclusive lineage-defining mutations with geographically restricted patterns of distribution has potential implications for diagnosis, pathogenesis and vaccine design. Our work provides novel insights on the temporal evolution of SARS-CoV-2.

**Importance:** The SARS-CoV-2 / COVID-19 pandemic has spread far and wide with high infectivity. However, the severeness of the infection as well as the mortality rates differ greatly across different geographic areas. Here we report high frequency mutations in the SARS-CoV-2 genomes which show the presence of linage-defining, leading and trailing mutations. Moreover, we propose for the first time, five mutually exclusive clusters of SARS-CoV-2 which account for 75% of the genomes analysed. This will have implications in diagnosis, pathogenesis and vaccine design

## Introduction

The COVID-19 pandemic caused by the novel coronavirus SARS-CoV-2 (SARS-like coronavirus 2) is rapidly spreading across the globe with over 10 million cases within a period of about 6 months (https://www.who.int/emergencies/diseases/novel-coronavirus-2019). This is far more alarming than the SARS (Severe Acute Respiratory Syndrome) epidemic of 2003 which affected 26 countries, killed 774 people, but was contained within 6 months (https://www.who.int/csr/sars/country/table2003_09_23/en/). Understanding the unique features of the SARS-CoV-2 is key to containing the COVID-19 pandemic (1). SARS-CoV and the SARS-CoV-2 are closely related RNA viruses with plus strand RNA genomes. RNA viruses usually display high mutation rates that facilitate adaptability and virulence (2). The moderate mutation rates seen in SARS-CoV (3) have been attributed to the presence of a 3’-5’ exonuclease or proof-reading like activity (4).

The rapid global spread of SARS-CoV-2 in a short period of time and the availability of a large number of fully sequenced genomes provide us with a unique opportunity of understanding the short-term temporal evolution of this virus in humans in a near real-time scale. While several studies have used phylogenetic analysis to reveal mutation hotspots in the viral genome and identify variants (5–7), we have utilized an alternative approach by focusing only on mutations present in >1% genomes followed by clustering. By this approach we propose the classification of the SARS-CoV-2 virus genomes into 5 mutually exclusive lineages with unique set of co-occurring mutations and geographic distribution.

## Results and discussion

### Mutation landscape of SARS-CoV-2

To understand the evolution of SARS-CoV-2 over the first three months of the pandemic, we performed a detailed analysis of full-length genomes available from the GISAID database from December 24, 2019 to March 24, 2020. A total of 1448 SARS-CoV-2 sequences (>29000 nt) were retrieved from GISAID (as on 24^th^ March 2020). A multiple sequence alignment was performed to visualize the variations in SARS-CoV-2 genomes. To analyse the single nucleotide substitutions in detail, we decided to focus on those which were observed in >1% of the genomes. As expected, the 5’ and 3’ ends of the sequences contained gaps and inconclusive nucleotide positions (Ns) attributable at least in part to the technical shortcomings of sequencing. The substitutions were considered only when the quality of the sequences at a given nucleotide position is greater than 95%. Our analysis revealed a total of 40 nucleotide substitutions which occurred at > 1% in the SARS-CoV-2 genomes (Table 1 and Figure 1A). This includes 15 synonymous mutations, 22 missense mutations and 3 substitutions in the non-coding regions. ORF9/N gene encoding for the nucleocapsid phosphoprotein seems to have accumulated the maximum number (n=8) of mutations. This is also evident when substitution frequencies for each gene are normalized for the gene length (Figure 1B). Among the non-structural proteins (Nsps), the viral RNA polymerase-helicase pair (nsp12/13) accumulated nine mutations. These observations are important considering that the widely used diagnostic tests for the detection of SARS-CoV-2 target nsp12 (RdRP, RNA-dependent RNA Polymerase) and N genes. While, we found 2 mutations in the Nsp3 region, the substitution frequencies per unit length are rather low as Nsp3 is the longest among the non-structural genes. It is interesting to note than no mutations were observed in the Nsp5 region coding for the main protease (Mpro), a widely investigated anti-viral target for SARS-CoV-2 (8, 9). A quick scan along the 30kb genome of SARS-CoV-2 clearly suggests clustering of mutations at the 3’ end of the genome (Figure 1). It would be worth investigating differences if any, in the fidelity of the virus polymerase or exonuclease pertaining to the 3’ end of the genome. All subsequent analysis presented in this study pertain to the 40 mutations occurring at >1% frequency.

**Figure 1.**
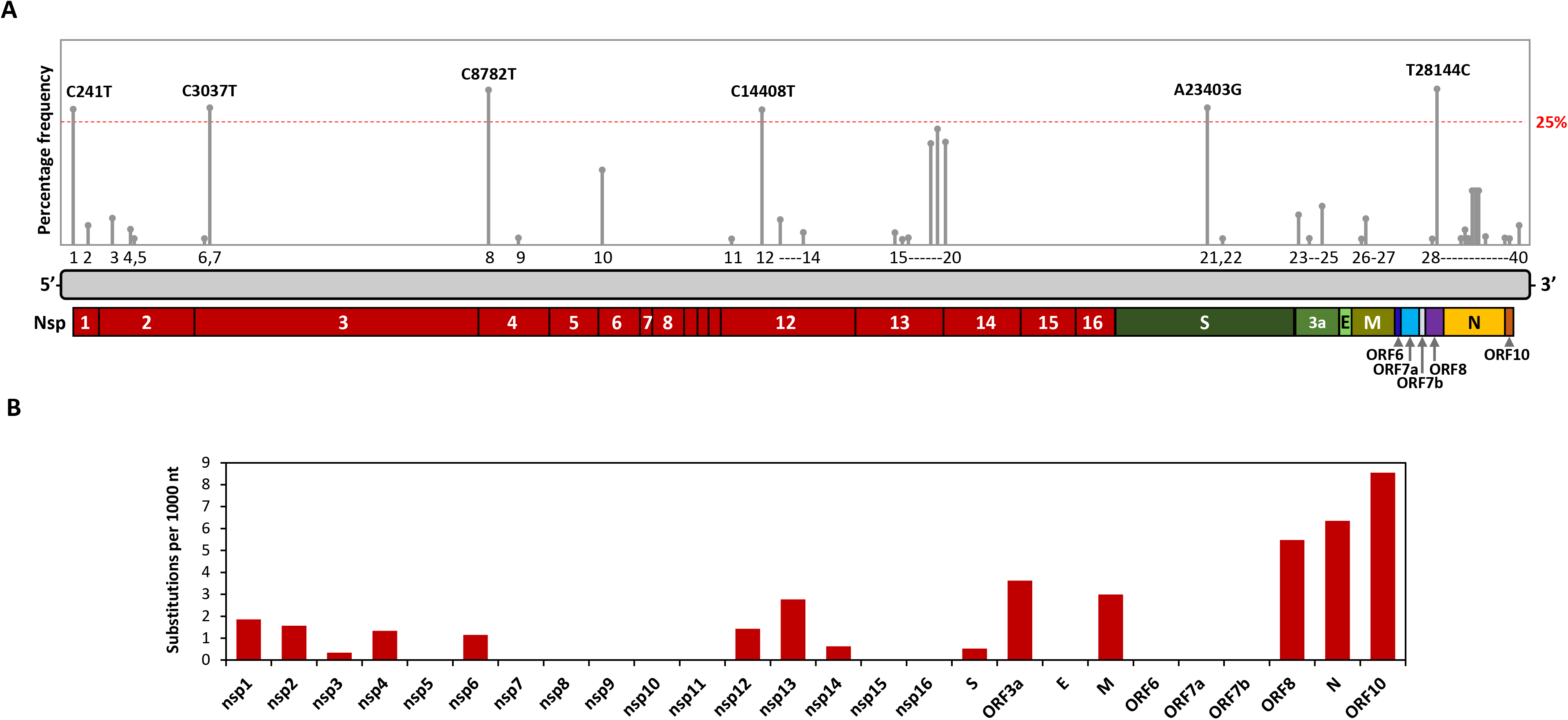
Mutation landscape of SARS-CoV-2. **A.** A schematic representation of the SARS-CoV-2 genome with gene/polypeptide annotations based on the NCBI reference sequence. The 40 substitution (>1%) sites are indicated with grey bars. The height of the bars indicates the percentage frequency of each mutation in the 1448 genome dataset and mutations that occur at >25% are labelled. **B.** Substitutions per unit length (1000 nt) per gene are calculated and plotted. While the N gene (8 substitutions in 1259 nt) and the ORDF10 displayed the highest number of substitutions per unit length, it should be noted that the putative ORF10 is only 117 nt long and harbours a single substitution.

**Table 1.**
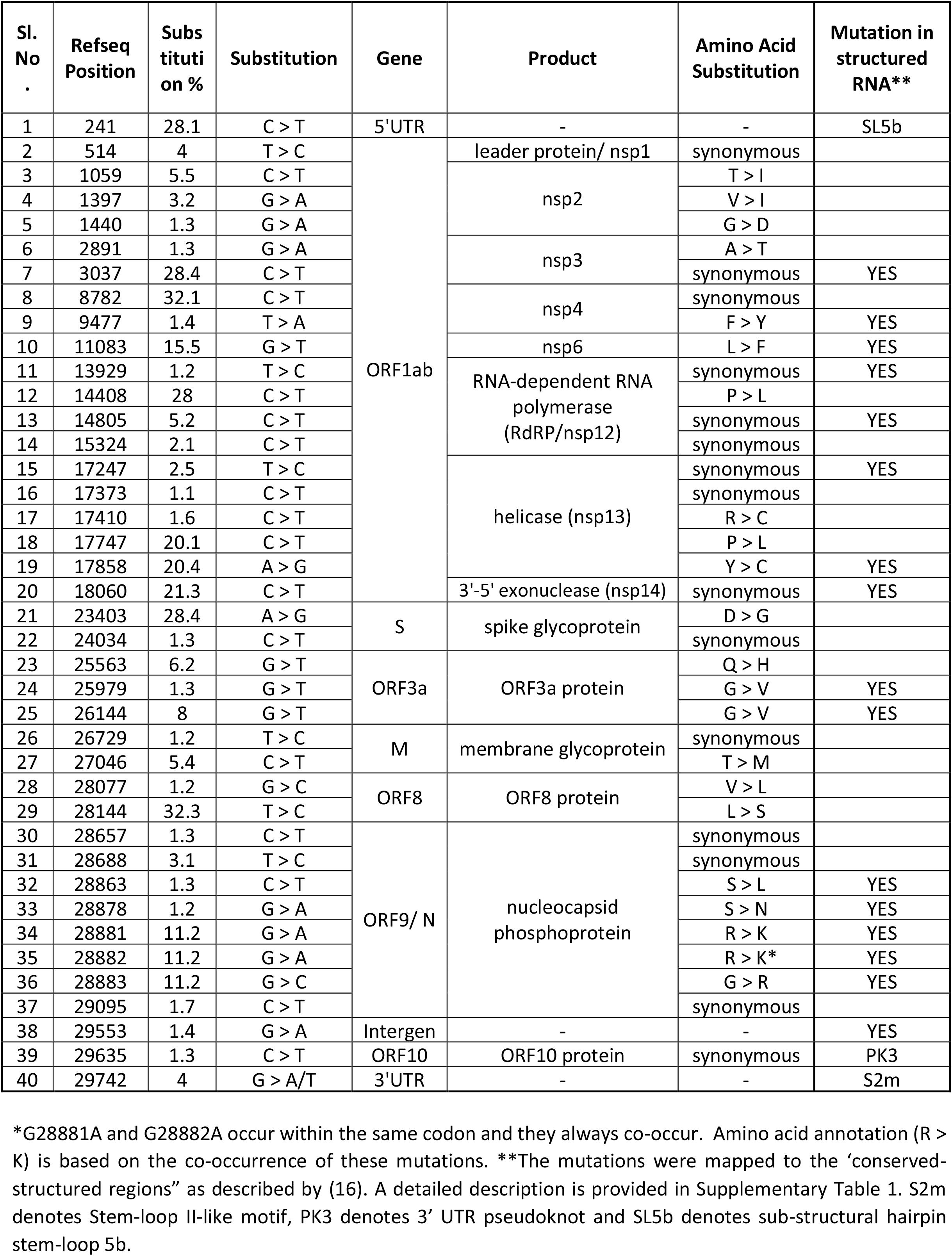
Summary of single nucleotide substitutions with >1% frequency in SARS-CoV-2 genomes.

### Mapping mutations to conserved structural elements of SARS-CoV-2 genome

The mutations identified in the non-coding regions include one each from the 5’ UTR (C241T), 3’ UTR (G29742A/T) and the intergenic region between ORF9 and ORF10 (G29553A) (Table 1). These mutations may have implications on viral RNA folding. Group I and II coronaviruses harbour three sub-structural hairpins SL5a, SL5b & SL5c with conserved hexameric “UUYCGU” loop motifs, which when present in multiple copies are presumed to act as packaging signals for viral encapsidation (10). Similar structures have been predicted in the 5’UTR of SARS-CoV-2 (11, 12). The C241T mutation changes the C-residue in the hexameric loop motif of SARS-CoV-2 SL5b (Figure 2A & 2B). This is one of the earliest and the most prevalent (28.1%) of the SARS-CoV-2 mutations and may impact viral packaging and titres.

**Figure 2.**
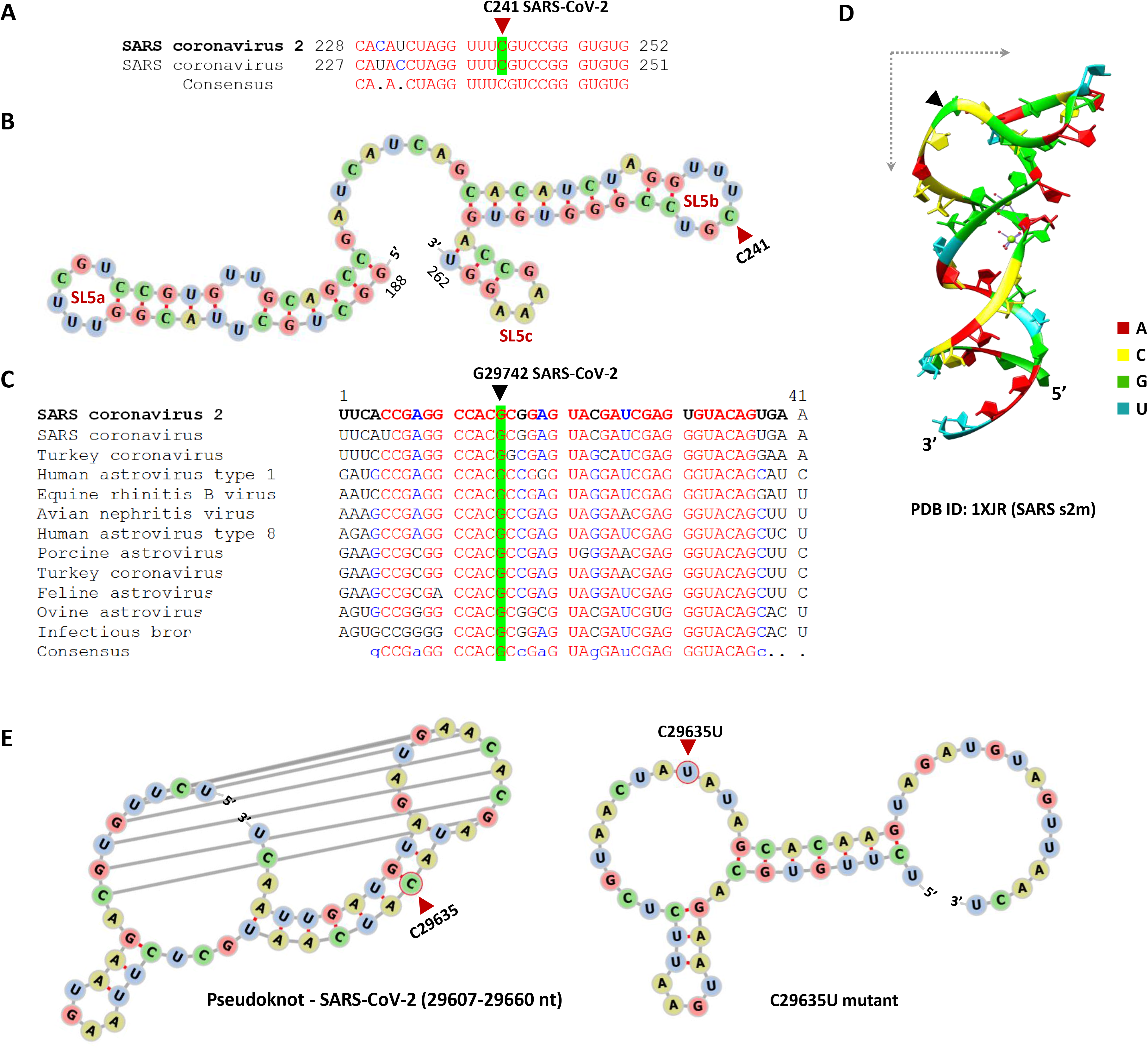
Structural elements in the SARS-CoV-2 and nucleotide substitutions. **A.** Alignment of the 5’ UTR SL5b stem-loop region from SARS-CoV and SARS-CoV2 genomes showing the conserved loop residue – C241 in SARS-CoV-2 which is mutated to T in 28.1% of genomes analysed. **B.** The 5’UTR stem-loops SL5a, SL5b and SL5c from SARS-CoV-2 depicting the position of C241 residue in the hexameric loop of SL5b. **C.** Multiple alignment of stem-loop II-like motif (s2m) in the 3’ UTR of coronaviruses with the G29742 mutation hotspot in SARS-CoV-2 highlighted. The SARS-CoV and SARS-CoV-2 sequences are nearly identical. **D.** The crystal structure of SARS-CoV s2m (PDB ID: 1XJR) showing the crucial position of the conserved G residue in the right-angle bent region of the stem-loop. **E.** The left panel shows the predicted structure of the 3’UTR pseudoknot (PK3) region in wild-type SARS-CoV-2 genome. The predicted structural changes associated with the C29635U mutant is shown in the right panel.

Interestingly, G29742 in the 3’ UTR is the only position among the 40 mutations which shows two different variants, G29742T and G29742A (Table 1). This residue is part of the stem-loop II-like motif (s2m) highly conserved in SARS-like coronaviruses (Figure 2C). SARS-CoV s2m sequence assumes a unique RNA fold similar to that seen in ribosomal RNA and is thought to be of regulatory function (13). The G29742 occupies a conserved position critical for the unique kinked structure and the substitutions at this position are expected to alter the s2m stem-loop (Figure 2D). Recent studies have also reported other low frequency mutations in this region (7). Another structural element relevant for coronavirus replication is the hairpin region designated as PK3 (3’ UTR pseudoknot)(14). While the PK3 stem-loops (29607-29660 nt in SARS-CoV-2) are present in the 3’ UTR of other coronaviruses, SARS-CoV-2 harbors the predicted open-reading frame ORF10 overlapping with this region. The C29635T mutation falls within the pseudoknot sequence and comparison of predicted structures for the wild-type and C29653T mutant PK3 display strong changes in the hairpin structure and orientation (Figure 2E). It is interesting to note that 17 out of 19 sequences harbouring this mutation have been reported from Japan in the month of February 2020.

Substitutions in the coding region may also have implications in viral RNA folding and structure (15). A recent study identified 106 conserved and highly structured RNA elements in the SARS-CoV-2 genome (16). We mapped eighteen of our 41 substitutions to these highly structured regions (Table 1 and Supplementary Table S1). We also aligned the SARS-CoV-2 genome with the genomes of closely related coronaviruses and analysed the positions homologous to these 40 nucleotide substitution sites (Supplementary Table S2). Interestingly, 12 substitutions resulted in nucleotide changes identical to those in one or more of the related coronavirus genomes.

### C to T and G to A substitutions predominate in the SARS-CoV-2 genomes

Transitions are more common than transversions across mammalian genomes as well as viruses (17–19). Analysis of substitutions in Influenza A virus and HIV-1 suggest that transversions are more deleterious than transitions (20). Hence, it is not surprising that 32 of the 40 substitutions we report here for SARS-CoV-2 are transitions. Substitutions at one of the nucleotide positions include both transition (G29742A) and transversion (G29742T). Thus the transitional mutation bias in SARS-CoV-2 is in keeping with that reported for other viruses (19).

C to T substitutions account for more than 40% (n=17) followed by G to A substitution which account for about 20% (n=8) of the substitutions (Figure 3A & 3B). The predominance of C to T (U) substitutions in SARS-CoV-2 has been documented recently (5, 21). A similar trend has also been reported for SARS-CoV (22). Amongst host RNA editing enzymes, APOBECs (Apolipoprotein B mRNA editing complex) and ADARs (Adenosine Deaminases Acting on RNA) have been studied for their ability to edit virus genomes (23, 24). Both negative-strand intermediates and genomic RNA (positive sense) may be available as ssRNA inside coronavirus infected host cells. The negative-strand intermediates serve as templates for the synthesis of the positive strand (genomic RNA) as well as the sub-genomic transcripts. In this process the negative-strand intermediates will progressively lose the ssRNA conformation and gain dsRNA conformation. In essence, both ssRNA and dsRNA forms will be available for RNA editing by host enzymes. This is important considering that RNA editing APOBECs specifically act on ssRNA to induce C>U transitions and ADARs target dsRNA resulting in A>I (inosine; read as G) editing. APOBEC-mediated editing of the genomic RNA (positive strand) will result in C to U changes, while the editing of the negative-strand intermediates (C to U) will result in G to A changes in the genomic RNA (positive strand). It is well documented that the abundance of the positive sense RNA outnumbers that of the negative strand RNA in infected host cell (25). Our finding of predominance of C to T (C to U) mutations (41.5%) over G to A mutations (19.5%) is consistent with the abundance of genomic RNA (positive sense) over the negative-strand intermediates.

**Figure 3.**
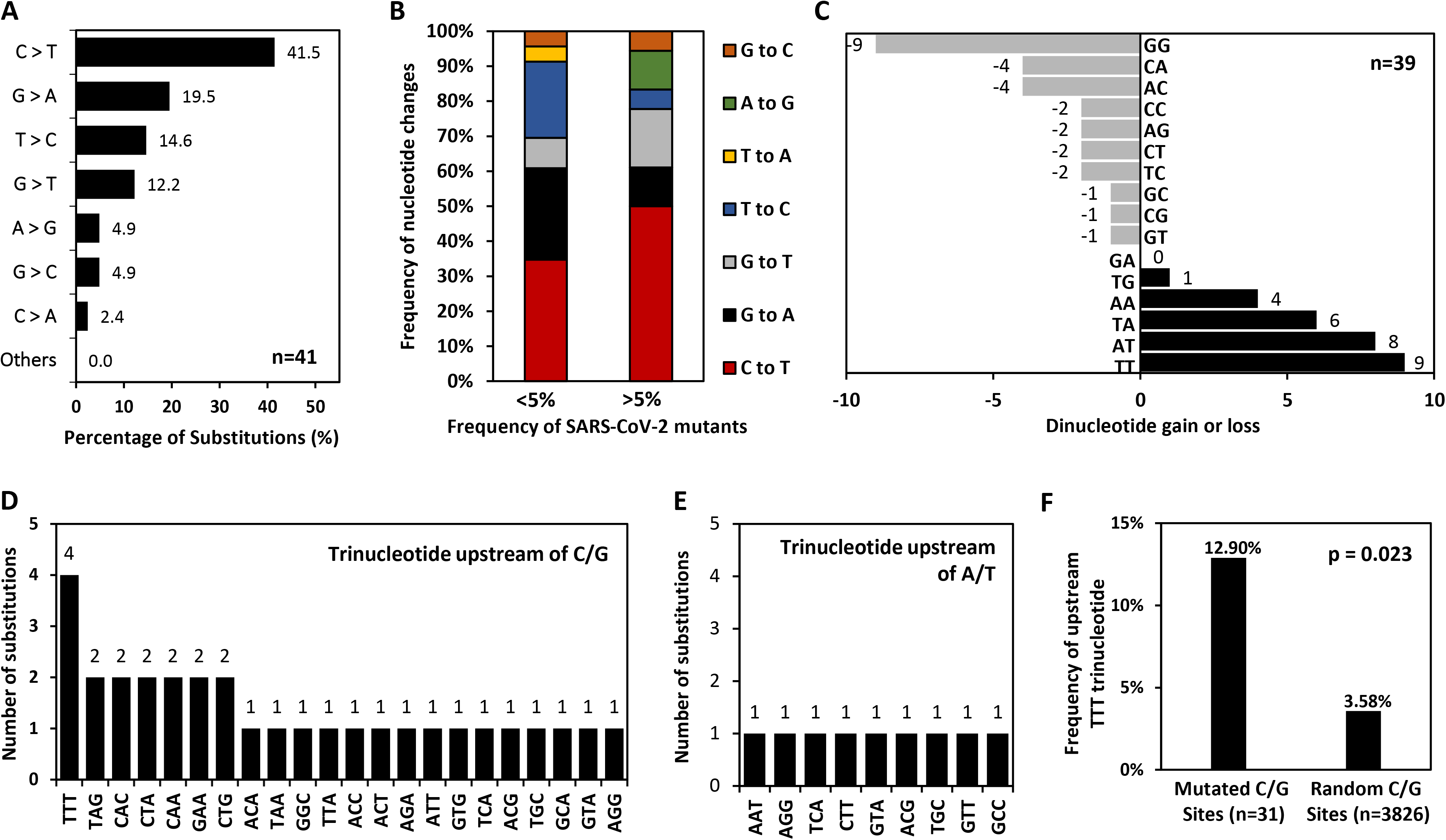
Nucleotide substitutions in the SARS-CoV-2 genomes are non-random. **A.** Percentage of specific nucleotide changes among the substitutions showing higher proportion of C>T and G>A changes. **B.** When mutations occurring at >5% (n=18) and <5% (n=23) are separately analysed, the distribution shows positive correlation between mutation frequency and enrichment of C>T substitutions. **C.** The net gain/loss of specific dinucleotides at the 39 single substitution sites (G29742T/A excluded) are calculated and plotted. **D & E.** The trinucleotide sequences upstream of C/G (D) and A/T (E) nucleotide substitution sites are shown. **F.** The frequency of appearance of TTT trinucleotides upstream of C/G substitution sites are compared to that of random C/G sites in the SARS-CoV-2 genome.

Similarly, ADAR-mediated editing of the genomic RNA (positive sense) will result in A to G changes and the editing of the negative-strand (A to G) will result in U to C changes in the genomic RNA (positive sense). Our findings indicate that the substitutions which are consistent with ADAR editing (A to G and U to C), constitute about 19.5 % (8 of 41 substitutions) of those observed in SARS-CoV-2 genomes analysed. The predominance of APOBEC-like RNA editing (i.e C to U and G to A) over ADAR-like RNA editing (i.e. A to G and U to C) in SARS-CoV-2 may be explained at least in part by the transient nature of virus dsRNA available inside infected host cells. Editing of SARS-CoV-2 RNA by APOBECs and ADARs has been recently reported (26). In addition, APOBEC3-mediated virus restriction has been documented for the human coronavirus HCoV-NL63 (27).

### Substitutions in SARS-CoV-2 lead to the loss of GG dinucleotides and gain of TT dinucleotides

We also analysed the dinucleotide context of the mutations described. For this purpose, we considered nucleotides flanking the substitutions described in table 1. For example, a C>T mutation in the trinucleotide sequence 5’-ACG-3’ sequence will result in 5’-ATG-3’ trinucleotide. In this context an AC and a CG dinucleotide are lost and an AT and a TG dinucleotide are gained. In other words, each substitution will lead to a loss of 2 dinucleotides and a gain of 2 dinucleotides. We analysed the 39 substitutions (G29742A/T was excluded) to assess the net loss or gain for each of the 16 dinucleotides. Interestingly, the 39 substitutions were associated with a loss of 9 GG dinucleotides and a gain of 9 TT dinucleotides (Figure 3C). The dinucleotides lost or gained are influenced not only by the nature of the substitutions (eg. C to T or G to A) but also by the nucleotides flanking the substitution site. Furthermore, dinucleotides provide important clues about virus pathogenesis (28, 29). In particular, CpG dinucleotides have been associated with virus replication, immune response and virus pathogenesis (30, 31). A recent paper suggests that SARS-CoV-2 is extremely CpG depleted to avoid host defences (32). Virus evolution has been also linked to variations in other dinucleotides including TpA and GpT (33, 34). Previous reports suggest that coronaviruses have excess of TT dinucleotides and are depleted for GG dinucleotides (35); this is consistent with major dinucleotide gain/loss in SARS-CoV-2 genomes (Figure 3C) The biological implications of these findings remain to be determined.

### Identification of TTTC and TTTG as hotspots for mutations in the SARS-CoV-2 genome

Signatures of APOBEC editing (C to T and G to A) dominate the substitutions observed in the SARS-CoV-2 genomes (Figure 3A). A recent study on RNA editing of SARS-CoV-2 genome shows evidence for APOBEC editing (26). Nonetheless, the specific motifs for APOBEC-mediated editing of virus genomes remain poorly understood. The presence of 17 C to T mutations and 8 G to A mutation among the 40 mutations we report gave us an opportunity to analyse preferences, if any for specific dinucleotide and trinucleotide motifs flanking these substitutions. While we found no such preferential motifs flanking the C to T or G to A substitution sites, we found interesting upstream trinucleotide motifs when all possible C (C>T, C>A and C>G) & G (G>A, G>T and G>C) substitution sites were considered together We found TTT trinucleotides upstream of 12.9 % of all substitutions (4 out of 31) occurring at C or G residues (C/Gs) (Figure 3D). We did not find any specific trinucleotide preference downstream to these sites (data not shown). There were no specific di or trinucleotide preferences upstream of substitutions occurring at A/Ts (Figure 3E).

Intrigued by the preference for TTT trinucleotide upstream of substitutions occurring at C/Gs, we analysed the statistical significance of this finding. Briefly, we generated 10000 random numbers corresponding to nucleotide positions in the SARS-CoV-2 genome and found 3826 positions with C/G. We then analysed the frequency of TTT trinucleotides upstream of these 3826 positions. The TTT trinucleotide frequency immediately upstream of substitutions originating at C/Gs is significantly higher than that upstream of randomly generated nucleotide positions (12.9 % vs 3.6 %; p<0.05; Figure 3F). This finding indicates that TTT trinucleotides followed by C/G (i.e. TTTC or TTTG) in the SARS-CoV-2 genome represent hotspots for substitutions. To the best of our knowledge, tetranucleotide motifs predisposed to higher substitution rates have not been reported for SARS-CoV-2. Furthermore, TTT trinucleotides were not detected upstream of any substitution occurring at As or Ts (n=9) (Figure 3E).

While TTTG/TTTC have not been reported as hotspots among viruses, TTTG has been reported as a hotspot for G to T substitutions in yeast deficient in nucleotide excision repair (36). TTC has been identified as a hotspot for APOBEC editing among gamma herpesviruses, which are dsDNA viruses (37). The specific mechanism associated with increased substitutions at TTTG/TTTC motifs in the SARS-CoV-2 genome is unclear. As negative-strand intermediates of SARS-CoV-2 serve as the template for the synthesis of positive-sense genomic RNA, we speculate that the virus RdRP (RNA-dependent RNA polymerase, nsp12) may be more error prone at C/Gs following a homopolymeric stretch of Ts (i.e. TTT). We cannot rule out a role for other proteins involved in genomic RNA replication including the nsp13 helicase and the nsp14 exonuclease in this process. Nonetheless, the identification of TTTG and TTTC as hotspots for substitutions opens up a plethora of opportunities for research on identification of the underlying virus /host factors.

The longest homopolymeric stretch in the entire SARS-COV-2 genome is an octamer of Ts (i.e.TTTTTTTT) and the G at the end of this octamer (i.e. TTTTTTTTG) is one of the substitution sites (G11083T; present in 15.5% of sequences analyzed)(refer Table 1). More interestingly, a homopolymeric stretch of GGGG (nt.28881 to 28884 in the Refseq) is associated with a triple mutant (i.e. GGGG to AACG; present in 11.2% of sequences analysed). All the three Gs are either mutated together or they remain wild-type in the 1448 genomes analysed. In addition, among the 40 substitutions we report, only the G28881A and the G28882A represent mutations within a single codon (R203K in the nucleocapsid protein) in SARS-CoV-2. Multiple substitutions within a single codon represent positive selection of an amino acid (38). Further, positively selected amino acids are frequently present in virus proteins involved in critical functions such as receptor binding (39).

### Nucleotide substitutions define several clusters of SARS-CoV-2 genomes

Several studies have looked into the aspect of real-time SARS-CoV-2 evolution and strain diversification by using phylogenetic analyses (5–7). In contrast to this approach, we utilized our catalogued set of single nucleotide substitutions to understand the emergence of SARS-CoV-2 variants. A clustering analysis was performed to systematically group all 1448 full-length genomes into subtypes which are defined by the set of mutations they harbor. After removing groups which consisted of <15 genomes (approximately 1% of genomes analysed), the analysis revealed 23 distinct clusters (Figure 4A). This analysis identified very interesting geographical distribution of the clusters across the globe (Figure 4B). For example, cluster-6 (C8782T, T28144C) primarily consisted of sequences from Asia (AS). Interestingly, cluster-1 consisting of three additional mutations (cluster-6 mutations + C17747T, A17858G and C18060T) accounted for about 20% of the genomes analysed and was almost exclusively reported from North America (NA). A similar pattern was observed for cluster 16, which was also restricted to NA. The cluster 5 consisting of 77 sequences with 8 mutations (C241T, C3037T, C14408T, A23403G, C27046T, G28881A, G28882A, G28883C) was almost exclusively found in Europe (EU). Similarly, cluster 10 with a single mutation (T514C) was also predominantly reported from Europe.

**Figure 4.**
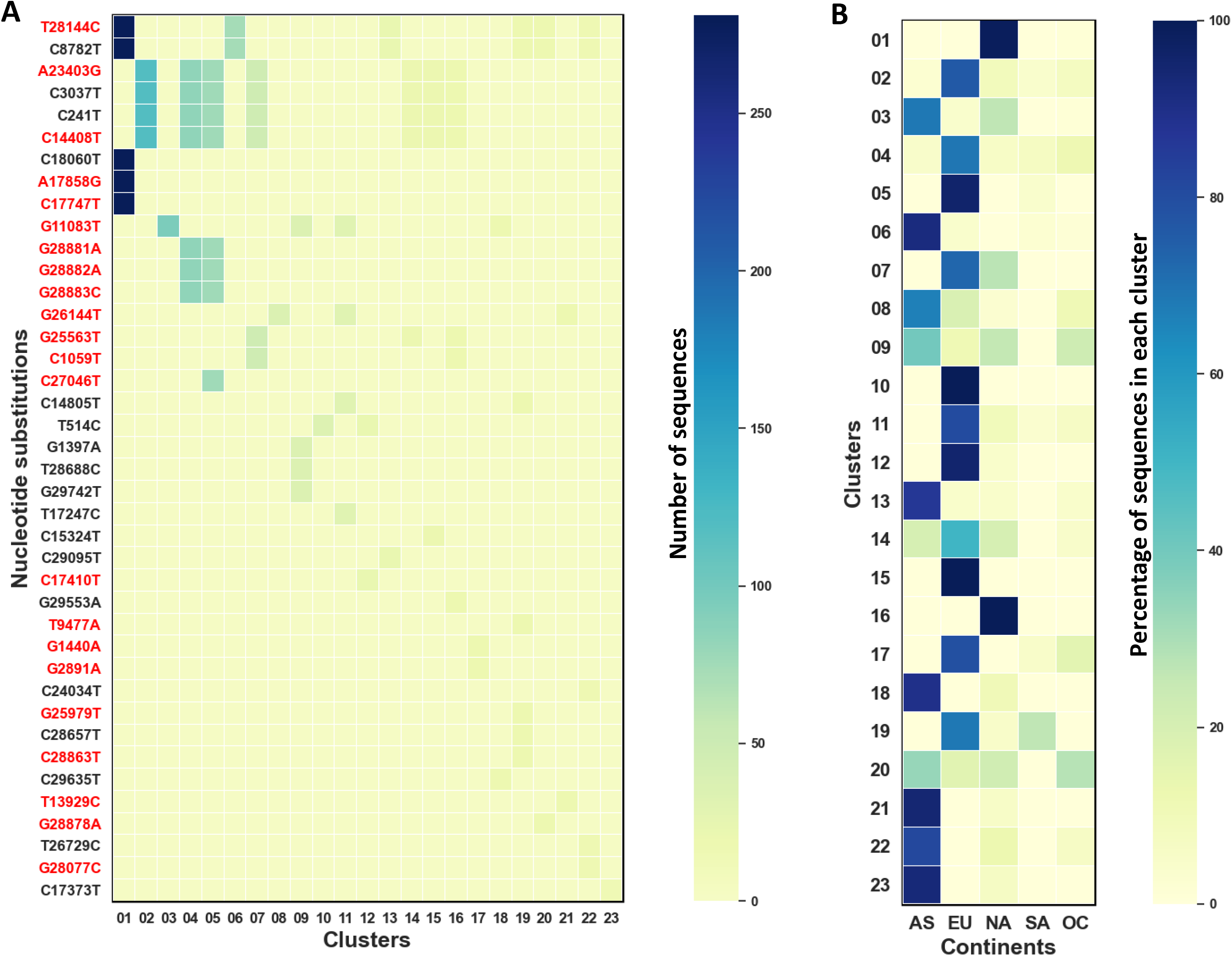
Clusters of SARS-CoV-2 genomes and their geographical distribution. **A.** Heat-map of the 23 clusters, generated from 1448 full-length genome sequences based on 40 single nucleotide substitutions. Color of the plot indicates the number of sequences with a given nucleotide substitutions present within the cluster. A specific mutation cluster is represented only if there are at least 15 full-length sequences. **B.** Heat-map showing the percentage distribution of clusters across different geographical regions (AS = Asia, EU= Europe, NA = North America, SA = South America and OC = Oceania).

### Identification of five mutually exclusive lineages and sequential mutation trails of SARS-CoV-2

Our clustering analysis revealed several mutations which almost always co-occur. We define co-occurring mutations as those which occur together in >95% of their individual occurrences. For example, if mutation A occurs 50 times and mutation B occurs 51 times in the dataset and mutations A and B are present together in the same genomes (i.e. co-occur) 50 times, they are defined as co-occurring mutations as 50/51 is >95%. Accumulation of mutations is key to the diversification of viral lineages. In addition, the co-occurrence of mutations in virus genomes is often suggestive of compensatory mutations. Individual mutations as well as co-occurring mutations can act as lineage defining mutations. We consider a specific mutation or a set of co-occurring mutations as “lineage-defining” for SARS-CoV-2, only when they are present in at least 2% (n=30) of the sequences analysed. Each lineage-defining mutation(s) are mutually exclusive and are not present along with another lineage-defining mutation. Our analyses reveal five mutually exclusive lineages for SARS-CoV-2. We refer to these five lineages as A1, B1, C1, D1 and E1in the chronological order of their appearance (Figure 5 and Supplementary Figure S1). These five lineages account for about 75% of the 1448 sequences. The A1 lineage (co-occurrence of C8782T, T28144C) appeared as early as 5^th^ January 2020 in Asia. The presence of three co-occurring mutations (G1397A, T28868C, G29742T) defines our B1 lineage. This lineage was also first observed in samples collected from Asia on 18^th^ January 2020. A single lineage defining mutation G26144T first reported from North America on the 22^nd^ January 2020 represents our C1 lineage. The co-occurrence of C241T, C3037T, C14408T, A23403G defines our lineage D1 that was first detected on 20^th^ February 2020 in Italy, Europe. The E1 lineage appeared later in The Netherlands (Europe) and is defined by the presence of the T514C mutation (Figure 4). The mutually exclusive lineages which we report here account for three quarters of all sequences and may facilitate the development of simple alternatives to whole genome sequencing for epidemiological typing. Off note, allelic discrimination assays for co-occurring single nucleotide variations have been successfully used for genotyping of SARS-CoV in the past (22, 40).

**Figure 5.**
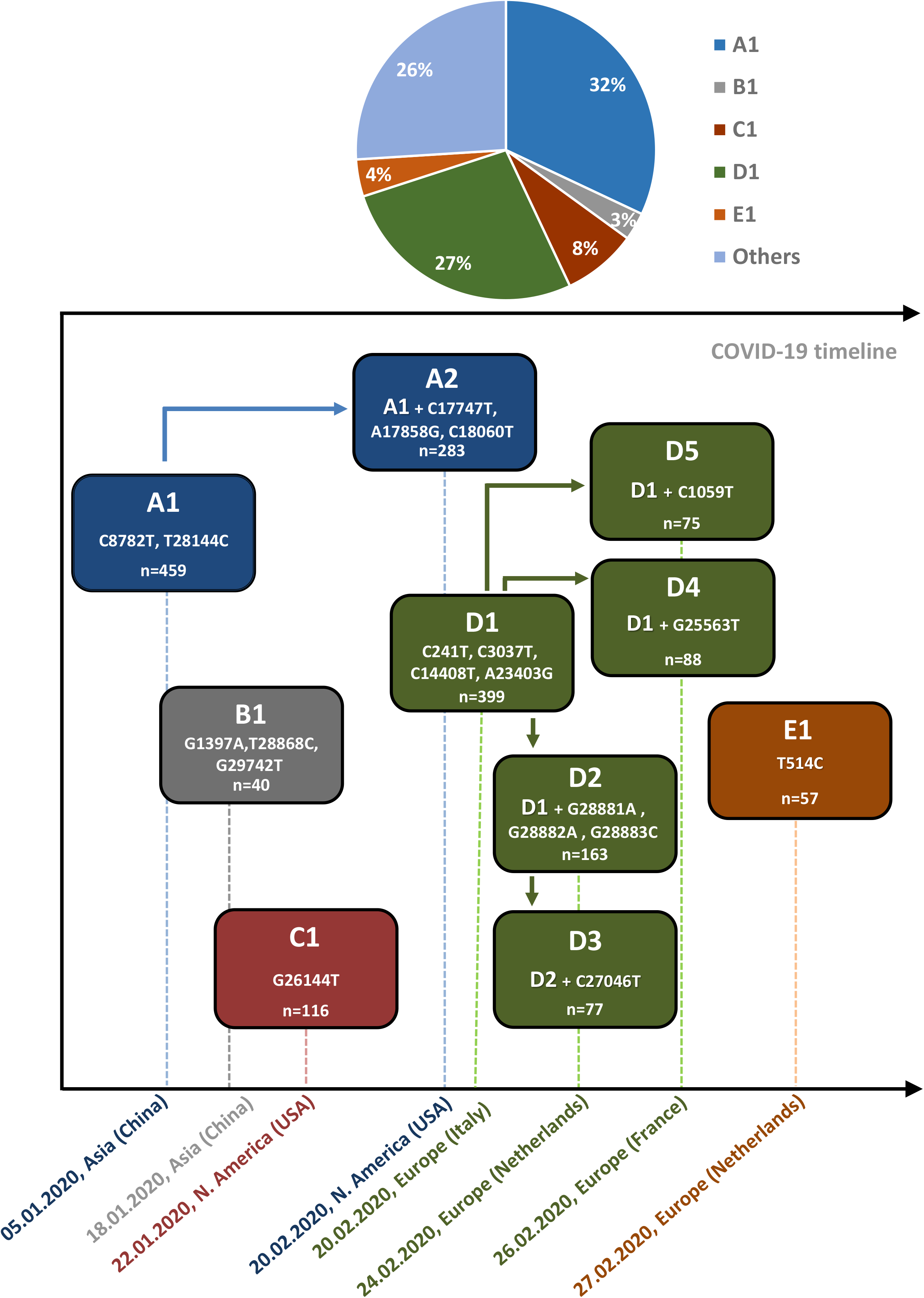
Five mutually exclusive lineages of SARS-CoV-2 and temporal acquisition of sequential mutations. Schematic representation of the COVID-19 pandemic time-line showing the leading mutations (A1, B1, C1, D1 and E1) and the sequential appearance of trailing mutations giving rise to additional SARS-CoV-2 variants (A2, and D2-D5). The date and country/continent of origin of sequences in which the indicated set of mutations were first reported are shown. The pie-chart in the upper panel shows the percentage distribution of the analysed genomes across the mutually exclusive lineages

Leading and trailing mutations have been previously described in virus genomes (41, 42). We define leading mutations as mutations that are documented in SARS-CoV-2 at an earlier time point and their presence is a pre-requisite for the origination of trailing mutations. In other words, trailing mutations do not occur in the absence of leading mutations. The lineage-defining mutations A1 through E1 represent leading mutations in the evolution of SARS-CoV-2. The chronology of the first appearance of all the 40 substitutions in the SARS-CoV-2 genome is summarised in Supplementary table S3. The A1 lineage accumulates a trailing mutation set giving rise to A2 (A1+ C17747T, A17858G, C18060T) variants of SARS-CoV-2. The A2 sub-lineage first appeared on 20^th^ February 2020 from the USA and accounts for about 20%% of the 1448 genomes. Even though the A1 lineage originated in Asia, the A2 sub-lineage (that is defined by 3 additional trailing mutations) appeared 6 weeks later in North America and remained restricted to this geographic region. (Supplementary Figure S2). Thus these three A2-defining trailing mutations can be used to discriminate between Asian and North American strains in the A1 lineage. Considering the fact that the A1 lineage accounts for >30% of the sequences analysed here, these observations may have potential implications in investigating possible intercontinental variations in infectivity and disease outcomes.

The most diverse pattern of trailing mutations has appeared in the D1 lineage, which incidentally is among the most recent of the lineage-defining mutations of SARS-CoV-2. Moreover, the lineage-defining mutations in the D1 variant include missense mutations in the Nsp12/RdRP (C14408T, P>L) and the S gene (A23403G, D>G), in addition to the SL5b loop mutation (C241T, Figure S1). While D2 (D1+G28881A, G28882A, G28883C), D4 (D1+G25563T) and D5 (D1+C1059T) emerged from D1 when the virus acquired specific set of trailing mutations, D3 (D2+C27046T) originated from the D2 variant upon the gain of a missense mutation (C27046T, T>M) in the viral M protein. Interestingly, the D1 lineage is mostly restricted to Europe and North America (Supplementary Figure S2) and involves incorporation of missense mutations in the viral N (D2 trailing mutations C28881A, G28882A, G28883C), M (D3 trailing mutation C27046T), ORF3a (D4 trailing mutation G25563T) and Nsp2 (D5 trailing mutation C1059T) genes (Figure 4 and Table 1). Within the D1 lineage, we could observe interesting bifurcation based on geographical distribution with D2 and D3 prevalent in Europe and D4 and D5 in Europe and North America. E1 (T514C, a synonymous mutation in the leader peptide/Nsp1), the most recent of the lineage-defining mutations of SARS-CoV-2 is predominantly seen in European strains (Supplementary Figure S2).

### Dynamics of single-nucleotide substitutions in the SARS-CoV-2 genome

The constantly updating list of SARS-CoV-2 sequences provides a unique opportunity for us to understand the evolution of this virus in humans in the course of the global outbreak. To understand the dynamics of different mutations over the evolution of the virus, we compared full-length genome datasets from GISAID with submission dates until 31^st^ January (n=210), 29^th^ February (n=664) and 24^th^ March 2020 (n= 1448). The mutations which show significant change in their frequency over the first three months of SARS-CoV-2 evolution are shown in Figure 6. The data is filtered for mutations which appeared in >5% of the sequences in the first (n=210) or complete (n=1448) dataset. Distinct patterns emerge for different mutations spread over the three datasets (Figure 6). Interestingly, the lineage-defining A1 mutation appear very early and despite slight reduction in the second dataset, consistently account for about one third of the sequences in each of the datasets. This would mean that the A1 lineage is consistently contributing to about a third of all newly reported sequences. The D1 lineage profile follows a logarithmic rise between the 2^nd^ and 3^rd^ datasets. The first case of this leading mutation appeared on the 20^th^ of February and by the 24^th^ of March it was among the most frequent mutations (399 out of 1448; >27%). Consistent with the appearance of the trailing mutations later in the timeline, trailing mutations for A2 (denoted as A2’), D2 (D2’), D3 (D3’), D4 (D4’) and D5 (D5’) sharply increase in numbers between the second and third dataset. C14805T (Nsp12/RNA polymerase synonymous) is another mutation which also show an upward tendency reaching 5.2% in the complete dataset (Figure 6A).

**Figure 6.**
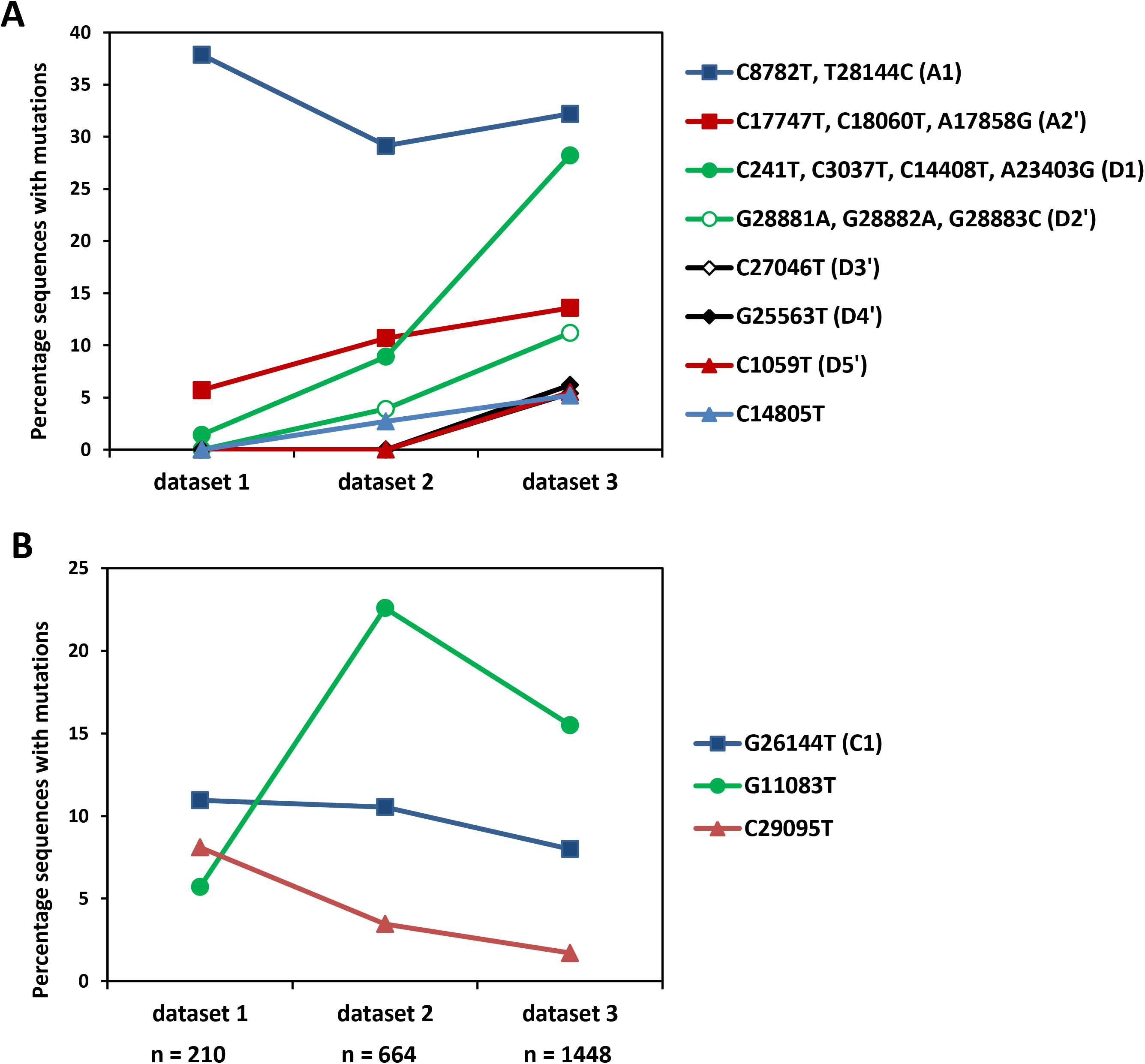
SARS-CoV-2 mutation frequencies over the first 3 months of the pandemic. We compared the percentage fraction of high frequency mutations (>5%) in three sequential datasets consisting of 210, 664 and 1448 genomes of SARS-CoV-2. **A.** Some mutations including the A1 and D1 leading mutations appear very early in the virus evolution and together they account for approximately 60% of the sequences in the most recent dataset. The trailing mutations of A2, D2, D3, D4, D5 are denoted as A2’, D2’, D3’, D4’, D5’ respectively and these do not include their leading mutations. For example, A2’ consists of A2 defining mutations alone and does not include its leading mutation set A1. A2’, D2’, D3’, D4’, D5’ as well as the C14805T mutations are increasing in frequency over time. **B.** C29095T and the C1 lineage mutation G26144T declined with time. The G11083T substitution shows an initial increase followed by a decrease, but retain significant presence in all three datasets.

The G11083T variant (Nsp6 L37F mutation) displays a profile similar to A1 between January and February, but shows a downward trend in march. However, this mutation is present in a significant proportion of sequences in the final dataset (15.5%) (Figure 6B). The C1 leading mutation shows a marginal downward trend in frequency over time. Interestingly, we also identified C29095T substitution which appear early in the timeline (Jan 2020) in >8% of the genomes, but their frequencies rapidly decline to <2% in the final dataset (Figure 6B).

Recent studies have shed light on a possible role for one of the D1 lineage-defining mutations (A23403G, Spike-glycoprotein D614G mutation) in the infectivity and fitness of SARS-CoV-2 (43). The D614G mutation has been linked to higher virus loads in patients which may explain the rapid spread of this variant. We have identified trailing mutations D2 through D5 associated with sub-lineages of the D614G variant (part of the D1 lineage). These findings provide the necessary groundwork for a plethora of studies with potential implications for diagnosis, pathogenesis and vaccine design.

## Conclusions

We focused our analyses on single nucleotide substitutions and identified 41 substitutions with >1% frequency in the SARS-CoV-2 genomes. Our analysis show that the nucleotide changes in the identified 41 substitution sites are non-random (predominant C to T/U and G to A) and are partially determined by the upstream sequence motifs. A significant proportion of analysed substitutions are consistent with APOBECs/ADARs editing. Clustering analysis revealed unique geographic distributions of SARS-CoV-2 variants defined by their mutation profile. Interestingly, we observed several co-occurring mutations that almost never occur individually. Our analyses reveal 5 mutually exclusive lineages of SARS-CoV-2 which account for about three quarters of the genomes analysed. Lineage-defining leading mutations in the SARS-CoV-2 genome precede the occurrence of trailing mutations that further define sub-lineages. The lineage-defining leading mutations together with the subsequently acquired trailing set of mutations provides a novel perspective on the temporal evolution of SARS-CoV-2

## Methods

### Sequence alignment and refinement of the SARS-CoV-2 genome sequences

The genomic Sequences of SARS-CoV-2 from clinical samples were downloaded from the GISAID database (https://www.gisaid.org/) (44). As of 24^th^ March 2020, GISAID consisted of 1448 SARS-CoV-2 full-length SARS-CoV-2 genomic sequences (>29000 nt) from human host. The sequences were downloaded at different time periods in the first three months of 2020 and fall into three serial datasets with submission dates until 31/01/2020 (n=210), 29/02/2020 (n=664) and 24/03/2020 (n=1448). The accession IDs of the genome sequences are provided as Supplementary table S4. The Alignment and refinement of the 1448 sequences with the SARS-CoV-2 reference genome were performed by using MUSCLE multiple sequence alignment software (45).. The genomic sequences for the bat coronavirus RaTG13 (Accession ID: MN996532.1), Pangolin corona virus (Accession ID: MT084071.1) and the SARS-CoV (Accession ID: NC_004718.3) were aligned with the SARS-CoV-2 reference genome to compare the substitution sites with homologous nucleotide positions in these closely related coronavirus genomes.

### Identification of single nucleotide substitutions in the SARS-CoV-2 genome

Alignment positions which harbor defined A/T/G/C residues (rather than gaps or ‘n’) in 95% of the aligned genomes were only considered for identifying nucleotide substitutions. These filtered nucleotide positions were analysed and scored for A/T/G/C occupancy in comparison to the SARS-CoV-2 reference genome (Accession: NC_045512.2). We then calculated the percentage of sequences with single-nucleotide variations for each nucleotide position in the SARS-CoV-2 genome. Substitutions with >1% frequency in the 1448 genome dataset were included for further analysis.

### Nucleotide, dinucleotide and trinucleotide variation analysis at substitution sites

The nucleotide changes at the 40 positions were analysed and tabulated. Similar analyses were done to calculate the gain and loss in frequencies of specific dinucleotide sequences at the substitution sites. We also analysed the trinucleotide context upstream and downstream of the substitution sites (−3 to +3 genomic positions). To understand the probability of finding TTT trinucleotide upstream of a random C/G position, we first mapped 10000 random positions (based on random numbers generated in MS Excel) on the SARS-CoV-2 genome and identified 3826 G/C residues. We the calculated the TTT trinucleotide frequency upstream to these 3826 sites and compared it to the actual frequencies at the substitution sites using a *chi*-squared test.

### Clustering analysis and defining leading and trailing mutations

Clustering was performed on the 1448 SARS-CoV-2 sequences with a custom script written using Python programming language and the data was visualized using Seaborn Statistical Visualization Tool (https://seaborn.pydata.org/). The 41 substitutions (39 substitutions + G29742T and G29742A) were used for the generation of mutation groups which co-occurred in many genomes. There were also sequences in which none of the selected mutations were present and hence formed the ‘no-mutation group’. In the second step, sequences harboring the same mutation groups were clustered together. Clusters with less than 15 genome sequences were excluded to obtain a list of 23 clusters represented as a heat map. The clusters were correlated with their geographical location (obtained from GISAID) and represented.

The datasets and clusters were analysed to identify co-occurring, lineage-defining, leading and trailing mutations. We classify mutations into the following categories:

- Co-occurring mutations: We define two (or more) mutations as co-occurring when they occur together in >90% of their individual occurrences.
- Leading and trailing mutations: we define leading mutations as mutations that are documented in SARS-CoV-2 genomes at an earlier time point and their presence is a pre-requisite for the development of trailing mutations. The trailing mutations never occur in the absence of the leading mutations. In view of possible sequencing errors, a maximum of 1 appearance of the trailing mutation(s) in the absence of the leading mutation(s) is tolerated.
- Lineage-defining mutations include both co-occurring mutations and singlet mutations that are present in ~2% (n=30) of sequences analysed. Lineage-defining mutations are mutually exclusive. To account for sequencing errors, a one sequence tolerance was allowed while deciding mutual exclusivity.

### RNA sequence alignment and structure analysis

RNA sequences were aligned using Multalin software (http://multalin.toulouse.inra.fr/multalin) (46). The stem-loop structures were predicted and visualized using the IPknot (47) (http://rtips.dna.bio.keio.ac.jp/ipknot/) and Forna web servers (48)(http://rna.tbi.univie.ac.at/forna/) respectively. The PDB structure for s2m stem-loop motif from SARS-CoV was downloaded from the Protein Data Bank (http://www.rcsb.org/pdb)(PDB ID: 1XJR) and was visualized using UCSF Chimera software (version 1.14)(49).

## Acknowledgements

We gratefully acknowledge the Authors from the Originating laboratories responsible for obtaining the specimens and the Submitting laboratories where genetic sequence data were generated and shared via the GISAID Initiative, on which this research is based (Supplementary Table 4). We acknowledge Kusuma School of Biological Sciences, Indian Institute of Technology Delhi for infrastructural support. The authors thank IIT Delhi HPC facility for computational resources.

## Legend to Supplementary Materials

**Supplementary Figure S1. The five lineage-defining mutations A1-E1 are mutually exclusive.** A checker board analysis showing the non-overlapping nature of the lineage-defining mutations A1 through E1. *The single exception was accession number EPI_ISL_414428 which had both D1 and E1 lineage defining mutations and was hence removed from both D1 and E1 lineages.

**Supplementary Figure S2. Geographical distribution of the five mutually exclusive lineages of SARS-CoV-2.** The pie diagrams represent the geographic distribution of the SARS-CoV2 lineages A1 (A1-A2), B1, C1, D1 (D1-D5) and E1.

**Supplementary Table S1. Substitutions in conserved-structured regions of SARS-CoV-2 genome.** The tabulated data show the mapping of the mutations to conserved-structured regions in SARS-CoV-2 genome. *The name and sequence interval of the conserved-structured region as described in Rangan et al., 2020 is shown.

**Supplementary Table S2. Nucleotide substitution sites in SARS-CoV-2 genome and homologous sites in related coronaviruses.** Multiple alignment of bat coronavirus RaTG13 (Accession ID: MN996532.1), Pangolin corona virus (Accession ID: MT084071.1) and the SARS-CoV (Accession ID: NC_004718.3) with the SARS-CoV-2 reference genome identified sites homologous to the nucleotide substitution sites. The positions showing differences from SARS-CoV-2 are listed here. The highlighted cells indicate positions with nucleotide-residues identical to the SARS-CoV-2 mutants. 25 of the 40 substitutions sites were conserved between the virus genomes. ‘-‘ indicate gaps in the sequence alignment.

**Supplementary Table S3. Chronological order of the first appearance of mutations in SARS-CoV-2.** The 41 substitutions/mutations (including G29742A and G29742T) in the SARS-CoV-2 genomes are shown with the date of collection and geographic location of the first reported sequence.

**Supplementary Table S4. Accession IDs of all SARS-CoV-2 sequences downloaded from GISAID including acknowledgement.** GISAID Accession IDs of all 1448 SARS-CoV-2 genome sequences used in the study with the acknowledgements and information on the participating labs.

## References

1. Wu A, Peng Y, Huang B, Ding X, Wang X, Niu P, Meng J, Zhu Z, Zhang Z, Wang J, Sheng J, Quan L, Xia Z, Tan W, Cheng G, Jiang T. 2020. Genome Composition and Divergence of the Novel Coronavirus (2019-nCoV) Originating in China. Cell Host Microbe 27:325–328.

2. Sanjuan R, Nebot MR, Chirico N, Mansky LM, Belshaw R. 2010. Viral mutation rates. J Virol 84:9733–48.

3. Zhao Z, Li H, Wu X, Zhong Y, Zhang K, Zhang YP, Boerwinkle E, Fu YX. 2004. Moderate mutation rate in the SARS coronavirus genome and its implications. BMC Evol Biol 4:21.

4. Minskaia E, Hertzig T, Gorbalenya AE, Campanacci V, Cambillau C, Canard B, Ziebuhr J. 2006. Discovery of an RNA virus 3’->5’ exoribonuclease that is critically involved in coronavirus RNA synthesis. Proc Natl Acad Sci U S A 103:5108–13.

5. Koyama T, Platt D, Parida L. 2020. Variant analysis of COVID-19 genomes. World Health Organ Preprint.

6. Forster P, Forster L, Renfrew C, Forster M. 2020. Phylogenetic network analysis of SARS-CoV-2 genomes. Proceedings of the National Academy of Sciences 117:9241–9243.

7. Yeh TY, Contreras GP. 2020. Faster de novo mutation of SARS-CoV-2 in shipboardquarantine. Bull World Health Organ Preprint.

8. Zhang L, Lin D, Sun X, Curth U, Drosten C, Sauerhering L, Becker S, Rox K, Hilgenfeld R. 2020. Crystal structure of SARS-CoV-2 main protease provides a basis for design of improved alpha-ketoamide inhibitors. Science 368:409–412.

9. Jin Z, Du X, Xu Y, Deng Y, Liu M, Zhao Y, Zhang B, Li X, Zhang L, Peng C, Duan Y, Yu J, Wang L, Yang K, Liu F, Jiang R, Yang X, You T, Liu X, Yang X, Bai F, Liu H, Liu X, Guddat LW, Xu W, Xiao G, Qin C, Shi Z, Jiang H, Rao Z, Yang H. 2020. Structure of M(pro) from COVID-19 virus and discovery of its inhibitors. Nature doi:10.1038/s41586/-020-2223-y.

10. Chen SC, Olsthoorn RC. 2010. Group-specific structural features of the 5’-proximal sequences of coronavirus genomic RNAs. Virology 401:29–41.

11. Chan JF-W, Kok K-H, Zhu Z, Chu H, To KK-W, Yuan S, Yuen K-Y. 2020. Genomic characterization of the 2019 novel human-pathogenic coronavirus isolated from a patient with atypical pneumonia after visiting Wuhan. Emerging Microbes & Infections 9:221–236.

12. Andrews RJ, Peterson JM, Haniff HF, Chen J, Williams C, Greffe M, Disney MD, Moss WN. 2020. An in silico map of the SARS-CoV-2 RNA Structurome. bioRxiv doi:10.1101/2020.04.17.045161:2020.04.17.045161.

13. Robertson MP, Igel H, Baertsch R, Haussler D, Ares MJr.,, Scott WG. 2005. The structure of a rigorously conserved RNA element within the SARS virus genome. PLoS Biol 3:e5.

14. Williams GD, Chang RY, Brian DA. 1999. A phylogenetically conserved hairpin-type 3’ untranslated region pseudoknot functions in coronavirus RNA replication. J Virol 73:8349–55.

15. Andrews RJ, Peterson JM, Haniff HS, Chen J, Williams C, Grefe M, Disney MD, Moss WN. 2020. An *in silico* map of the SARS-CoV-2 RNA Structurome. bioRxiv doi:10.1101/2020.04.17.045161:2020.04.17.045161.

16. Rangan R, Zheludev IN, Das R. 2020. RNA genome conservation and secondary structure in SARS-CoV-2 and SARS-related viruses: a first look. RNA doi:10.1261/rna.076141.120.

17. Rosenberg MS, Subramanian S, Kumar S. 2003. Patterns of transitional mutation biases within and among mammalian genomes. Mol Biol Evol 20:988–93.

18. Petrov DA, Hartl DL. 1999. Patterns of nucleotide substitution in Drosophila and mammalian genomes. Proc Natl Acad Sci U S A 96:1475–9.

19. Duchene S, Ho SY, Holmes EC. 2015. Declining transition/transversion ratios through time reveal limitations to the accuracy of nucleotide substitution models. BMC Evol Biol 15:36.

20. Lyons DM, Lauring AS. 2017. Evidence for the Selective Basis of Transition-to-Transversion Substitution Bias in Two RNA Viruses. Mol Biol Evol 34:3205–3215.

21. Wang C, Liu Z, Chen Z, Huang X, Xu M, He T, Zhang Z. 2020. The establishment of reference sequence for SARS-CoV-2 and variation analysis. J Med Virol doi:10.1002/jmv.25762.

22. Pavlović-Lažetić GM, Mitić NS, Tomović AM, Pavlović MD, Beljanski MV. 2005. SARS-CoV Genome Polymorphism: A Bioinformatics Study. Genomics, Proteomics & Bioinformatics 3:18–35.

23. Piontkivska H, Frederick M, Miyamoto MM, Wayne ML. 2017. RNA editing by the host ADAR system affects the molecular evolution of the Zika virus. Ecol Evol 7:4475–4485.

24. Cheng AZ, Yockteng-Melgar J, Jarvis MC, Malik-Soni N, Borozan I, Carpenter MA, McCann JL, Ebrahimi D, Shaban NM, Marcon E, Greenblatt J, Brown WL, Frappier L, Harris RS. 2019. Epstein-Barr virus BORF2 inhibits cellular APOBEC3B to preserve viral genome integrity. Nat Microbiol 4:78–88.

25. Sethna PB, Hofmann MA, Brian DA. 1991. Minus-strand copies of replicating coronavirus mRNAs contain antileaders. J Virol 65:320–5.

26. Di Giorgio S, Martignano F, Torcia MG, Mattiuz G, Conticello SG. 2020. Evidence for RNA editing in the transcriptome of 2019 Novel Coronavirus. bioRxiv doi:10.1101/2020.03.02.973255:2020.03.02.973255.

27. Milewska A, Kindler E, Vkovski P, Zeglen S, Ochman M, Thiel V, Rajfur Z, Pyrc K. 2018. APOBEC3-mediated restriction of RNA virus replication. Sci Rep 8:5960.

28. Wasson MK, Borkakoti J, Kumar A, Biswas B, Vivekanandan P. 2017. The CpG dinucleotide content of the HIV-1 envelope gene may predict disease progression. Sci Rep 7:8162.

29. Sankar S, Borkakoti J, Ramamurthy M, Nandagopal B, Vivekanandan P, Gopalan S. 2018. Identification of tell-tale patterns in the 3’ non-coding region of hantaviruses that distinguish HCPS-causing hantaviruses from HFRS-causing hantaviruses. Emerg Microbes Infect 7:32.

30. Simmonds P, Tulloch F, Evans DJ, Ryan MD. 2015. Attenuation of dengue (and other RNA viruses) with codon pair recoding can be explained by increased CpG/UpA dinucleotide frequencies. Proc Natl Acad Sci U S A 112:E3633–4.

31. Fros JJ, Dietrich I, Alshaikhahmed K, Passchier TC, Evans DJ, Simmonds P. 2017. CpG and UpA dinucleotides in both coding and non-coding regions of echovirus 7 inhibit replication initiation post-entry. Elife 6.

32. Xia X. 2020. Extreme genomic CpG deficiency in SARS-CoV-2 and evasion of host antiviral defense. Mol Biol Evol doi:10.1093/molbev/msaa094.

33. Upadhyay M, Samal J, Kandpal M, Vasaikar S, Biswas B, Gomes J, Vivekanandan P. 2013. CpG Dinucleotide Frequencies Reveal the Role of Host Methylation Capabilities in Parvovirus Evolution. Journal of Virology 87:13816–13824.

34. Simmonds P, Xia W, Baillie JK, McKinnon K. 2013. Modelling mutational and selection pressures on dinucleotides in eukaryotic phyla ‒selection against CpG and UpA in cytoplasmically expressed RNA and in RNA viruses. BMC Genomics 14:610.

35. Cheng X, Virk N, Chen W, Ji S, Ji S, Sun Y, Wu X. 2013. CpG usage in RNA viruses: data and hypotheses. PLoS One 8:e74109.

36. Abdulovic AL, Minesinger BK, Jinks-Robertson S. 2008. The effect of sequence context on spontaneous Polzeta-dependent mutagenesis in Saccharomyces cerevisiae. Nucleic Acids Res 36:2082–93.

37. Martinez T, Shapiro M, Bhaduri-McIntosh S, MacCarthy T. 2019. Evolutionary effects of the AID/APOBEC family of mutagenic enzymes on human gamma-herpesviruses. Virus Evol 5:vey040.

38. Bazykin GA, Dushoff J, Levin SA, Kondrashov AS. 2006. Bursts of nonsynonymous substitutions in HIV-1 evolution reveal instances of positive selection at conservative protein sites. Proc Natl Acad Sci U S A 103:19396–401.

39. Bush RM, Fitch WM, Bender CA, Cox NJ. 1999. Positive selection on the H3 hemagglutinin gene of human influenza virus A. Mol Biol Evol 16:1457–65.

40. Chung GTY, Chiu RWK, Cheung JLK, Jin Y, Chim SSC, Chan PKS, Lo YMD. 2005. A simple and rapid approach for screening of SARS-coronavirus genotypes: an evaluation study. BMC Infectious Diseases 5:87.

41. Neverov AD, Kryazhimskiy S, Plotkin JB, Bazykin GA. 2015. Coordinated Evolution of Influenza A Surface Proteins. PLoS Genet 11:e1005404.

42. Kryazhimskiy S, Dushoff J, Bazykin GA, Plotkin JB. 2011. Prevalence of Epistasis in the Evolution of Influenza A Surface Proteins. PLOS Genetics 7:e1001301.

43. Korber B, Fischer WM, Gnanakaran S, Yoon H, Theiler J, Abfalterer W, Hengartner N, Giorgi EE, Bhattacharya T, Foley B, Hastie KM, Parker MD, Partridge DG, Evans CM, Freeman TM, de Silva TI, McDanal C, Perez LG, Tang H, Moon-Walker A, Whelan SP, LaBranche CC, Saphire EO, Montefiori DC, Angyal A, Brown RL, Carrilero L, Green LR, Groves DC, Johnson KJ, Keeley AJ, Lindsey BB, Parsons PJ, Raza M, Rowland-Jones S, Smith N, Tucker RM, Wang D, Wyles MD. 2020. Tracking changes in SARS-CoV-2 Spike: evidence that D614G increases infectivity of the COVID-19 virus. Cell doi:https://doi.org/10.1016/j.cell.2020.06.043.

44. Elbe S, Buckland-Merrett G. 2017. Data, disease and diplomacy: GISAID’s innovative contribution to global health. Glob Chall 1:33–46.

45. Edgar RC. 2004. MUSCLE: a multiple sequence alignment method with reduced time and space complexity. BMC Bioinformatics 5:113.

46. Corpet F. 1988. Multiple sequence alignment with hierarchical clustering. Nucleic Acids Res 16:10881–90.

47. Kato Y, Sato K, Asai K, Akutsu T. 2012. Rtips: fast and accurate tools for RNA 2D structure prediction using integer programming. Nucleic Acids Res 40:W29–34.

48. Kerpedjiev P, Hammer S, Hofacker IL. 2015. Forna (force-directed RNA): Simple and effective online RNA secondary structure diagrams. Bioinformatics 31:3377–9.

49. Pettersen EF, Goddard TD, Huang CC, Couch GS, Greenblatt DM, Meng EC, Ferrin TE. 2004. UCSF Chimera--a visualization system for exploratory research and analysis. J Comput Chem 25:1605–12.

