## Supplementary Figure S1 for "Mutation landscape of SARS-CoV-2 reveals five mutually exclusive clusters of leading and trailing single nucleotide substitutions"

| Lineages | A1 (n=459) | B1 (n=40) | C1 (n=116) | D1 (n=400) | E1 (n=58) |
|---|---|---|---|---|---|
| A1 (n=459) | | 0 | 0 | 0 | 0 |
| B1 (n=40) | | | 0 | 0 | 0 |
| C1 (n=116) | | | | 0 | 0 |
| D1 (n=400) | | | | | 1* |
| E1 (n=58) | | | | | |

**Supplementary Figure S1. The five lineage-defining mutations A1-E1 are mutually exclusive.** A checker board analysis showing the non-overlapping nature of the lineage-defining mutations A1 through E1. *The single exception was accession number EPI_ISL_414428 which had both D1 and E1 lineage defining mutations and was hence removed from both D1 and E1 lineages.
