## Supplementary Figure S2 for "Mutation landscape of SARS-CoV-2 reveals five mutually exclusive clusters of leading and trailing single nucleotide substitutions"

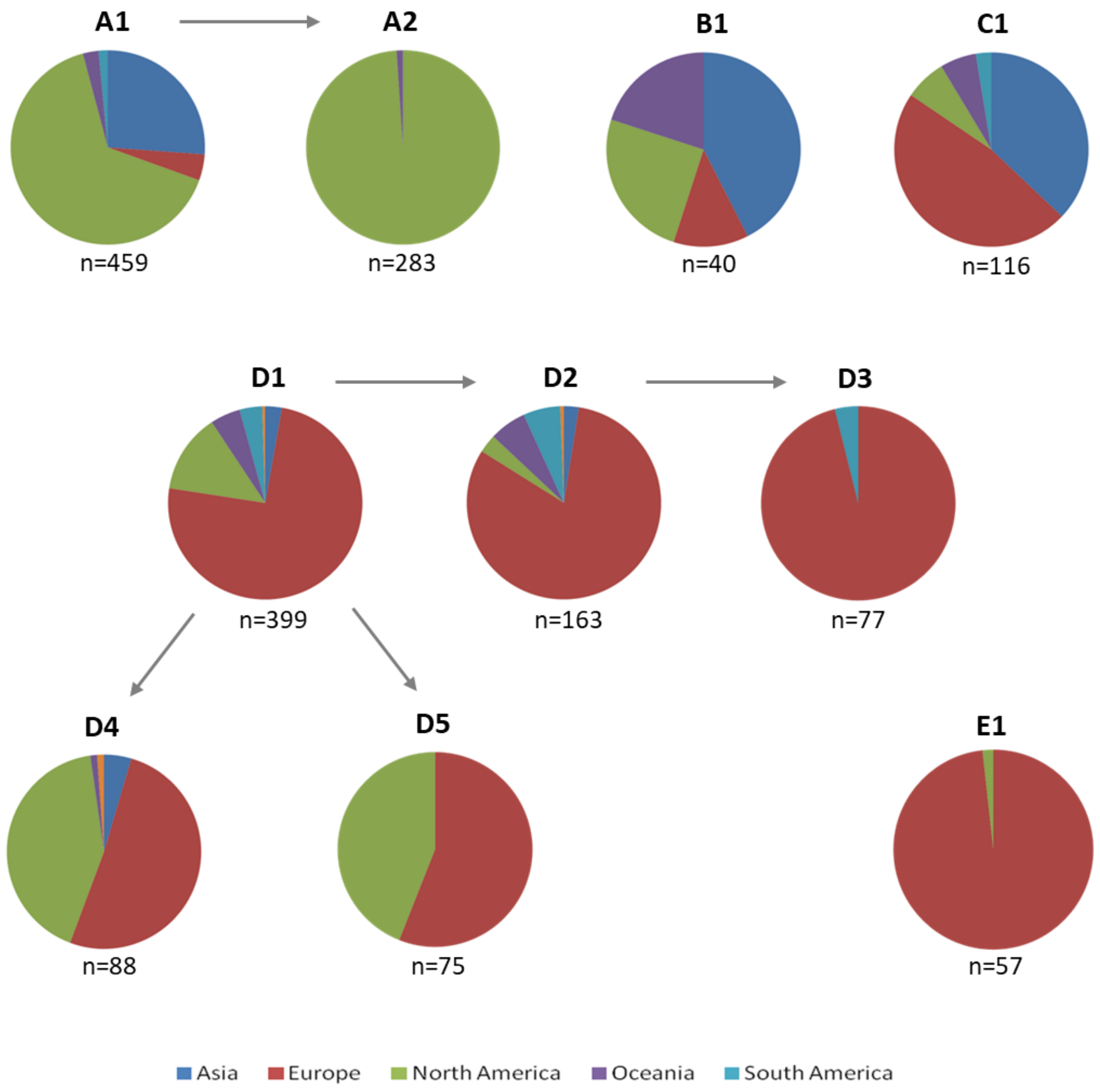


**Supplementary Figure S2. Geographical distribution of the five mutually exclusive lineages of SARS-CoV-2.** The pie diagrams represent the geographic distribution of the SARS-CoV2 lineages A1 (A1-A2), B1, C1, D1 (D1-D5) and E1.
