## Supplementary Table S1 for "Mutation landscape of SARS-CoV-2 reveals five mutually exclusive clusters of leading and trailing single nucleotide substitutions"

| Nucleotide Substitution | Gene/polypeptide | Substitution | Structured region* | |
|---|---|---|---|---|
| | | | Name | Sequence Interval |
| C3037T | nsp3 | synonymous | SARS-CoV-2-conservedstructured-19 | 2943-3062 |
| T9477A | nsp4 | Missense | SARS-CoV-2-conservedstructured-79 | 9410-9529 |
| G11083T | nsp6 | Missense | SARS-CoV-2-conservedstructured-29 | 10970-11089 |
| T13929C | ORF1ab | synonymous | SARS-CoV-2-conservedstructured-35 | 13886-14005 |
| C14805T | nsp12 | synonymous | SARS-CoV-2-conservedstructured-20, 74 | 14726-14845, 14766-14885 |
| T17247C | helicase (nsp13) | synonymous | SARS-CoV-2-conservedstructured-72 | 17205-17324 |
| C17410T | helicase (nsp13) | Missense | SARS-CoV-2-conservedstructured-54 | 17445-17564 |
| A17858G | helicase (nsp13) | Missense | SARS-CoV-2-conservedstructured-105 | 17765-17884 |
| C18060T | 3' -to-5' exonuclease | synonymous | SARS-CoV-2-conservedstructured-68 | 17965-18084 |
| G25979T | ORF3a | Missense | SARS-CoV-2-conservedstructured-10 | 25895-26014 |
| G26144T | ORF3a | Missense | SARS-CoV-2-conservedstructured-31 | 26095-26214 |
| C28863T | ORF9 / N | Missense | SARS-CoV-2-conservedstructured-104 | 28758-28877 |
| C28863T, G28878A, G28881A, G28882A, G28883C | ORF9 / N | Missense | SARS-CoV-2-conservedstructured-30 | 28838-28957 |
| G29553A | Intergenic | Non-coding | SARS-CoV-2-conservedstructured-33 | 29550-29669 |
| C29635T | ORF10 | synonymous | SARS-CoV-2-conservedstructured-33 | 29550-29669 |

**Supplementary Table S1. Substitutions in conserved-structured regions of SARS-CoV-2 genome.** The tabulated data show the mapping of the mutations to conserved-structured regions in SARS-CoV-2 genome. *The name and sequence interval of the conserved-structured region as described in Rangan et al., 2020 is shown.
