## Supplementary Table S2 for "Mutation landscape of SARS-CoV-2 reveals five mutually exclusive clusters of leading and trailing single nucleotide substitutions"

| Sl. No. | Substitution | SARS-CoV-2 | SARS-CoV | Bat -CoV | Pangolin-CoV |
|---|---|---|---|---|---|
| 1 | T514C | T | C | T | T |
| 2 | G1397A | G | A | G | A |
| 3 | G1440A | G | A | G | G |
| 4 | C3037T | C | T | T | C |
| 5 | C8782T | C | C | T | T |
| 6 | G11083T | G | T | G | - |
| 7 | C14805T | C | T | C | C |
| 8 | T17247C | T | T | T | C |
| 9 | C17373T | C | T | T | T |
| 10 | C18060T | C | T | T | A |
| 11 | C24034T | C | T | T | C |
| 12 | G25563T | G | A | G | - |
| 13 | C28657T | C | A | C | A |
| 14 | C29095T | C | T | T | T |
| 15 | C29635T | C | A | C | C |

**Supplementary Table S2. Nucleotide substitution sites in SARS-CoV-2 genome and homologous sites in related coronaviruses.** Multiple alignment of bat coronavirus RaTG13 (Accession ID: MN996532.1), Pangolin corona virus (Accession ID: MT084071.1) and the SARS-CoV (Accession ID: NC_004718.3) with the SARS-CoV-2 reference genome identified sites homologous to the nucleotide substitution sites. The positions showing differences from SARS-CoV-2 are listed here. The highlighted cells indicate positions with nucleotide-residues identical to the SARS-CoV-2 mutants. 25 of the 40 substitutions sites were conserved between the virus genomes. '-' indicate gaps in the sequence alignment.
