## Supplementary Table S3 for "Mutation landscape of SARS-CoV-2 reveals five mutually exclusive clusters of leading and trailing single nucleotide substitutions"

| Sl. No. | Mutation | % | Accession number | Date of first appearance | Geography |
|---|---|---|---|---|---|
| 1 | C241T | 28.1 | EPI_ISL_416327 | 28-Jan-2020 | >hCoV-19/Shanghai/SH0014/2020 |
| 2 | T514C | 4 | EPI_ISL_413586 | 27-Feb-2020 | >hCoV-19/Netherlands/Tilburg_1363354/2020 |
| 3 | C1059T | 5.5 | EPI_ISL_414625 | 26-Feb-2020 | >hCoV-19/France/PL1643/2020 |
| 4 | G1397A | 3.2 | EPI_ISL_412981 | 18-Jan-2020 | >hCoV-19/Wuhan/HBCDC-HB-05/2020 |
| 5 | G1440A | 1.3 | EPI_ISL_414497 | 25-Feb-2020 | >hCoV-19/Germany/NRW-02-1/2020 |
| 6 | G2891A | 1.3 | EPI_ISL_414497 | 25-Feb-2020 | >hCoV-19/Germany/NRW-02-1/2020 |
| 7 | C3037T | 28.4 | EPI_ISL_416327 | 28-Jan-2020 | >hCoV-19/Shanghai/SH0014/2020 |
| 8 | C8782T | 32.1 | EPI_ISL_406801 | 5-Jan-2020 | >hCoV-19/Wuhan/WH04/2020 |
| 9 | T9477A | 1.4 | EPI_ISL_416372 | 1-Feb-2020 | >hCoV-19/Shanghai/SH0069/2020 |
| 10 | G11083T | 15.5 | EPI_ISL_408480 | 17-Jan-2020 | >hCoV-19/Yunnan/IVDC-YN-003/2020 |
| 11 | T13929C | 1.2 | EPI_ISL_412029 | 30-Jan-2020 | >hCoV-19/Hong_Kong/VB20024950-2/2020 |
| | | | EPI_ISL_417183 | | >hCoV-19/Hong_Kong/HKPU23_2601/2020 |
| 12 | C14408T | 28 | EPI_ISL_412973 | 20-Feb-2020 | >hCoV-19/Italy/CDG1/2020 |
| 13 | C14805T | 5.2 | EPI_ISL_412116 | 9-Feb-2020 | >hCoV-19/England/09c/2020 |
| 14 | C15324T | 2.1 | EPI_ISL_406533 | 22-Jan-2020 | >hCoV-19/Guangzhou/20SF206/2020 |
| 15 | T17247C | 2.5 | EPI_ISL_413019 | 26-Feb-2020 | >hCoV-19/Switzerland/1000477102/2020 |
| | | | EPI_ISL_414011 | | >hCoV-19/England/200990006/2020 |
| 16 | C17373T | 1.1 | EPI_ISL_406535 | 22-Jan-2020 | >hCoV-19/Foshan/20SF210/2020 |
| | | | EPI_ISL_406536 | | >hCoV-19/Foshan/20SF211/2020 |
| | | | EPI_ISL_411066 | | >hCoV-19/Fujian/13/2020 |
| 17 | C17410T | 1.6 | EPI_ISL_413586 | 27-Feb-2020 | >hCoV-19/Netherlands/Tilburg_1363354/2020 |
| 18 | C17747T | 20.1 | EPI_ISL_413456 | 20-Feb-2020 | >hCoV-19/USA/WA-S2/2020 |
| 19 | A17858G | 20.4 | EPI_ISL_413456 | 20-Feb-2020 | >hCoV-19/USA/WA-S2/2020 |
| 20 | C18060T | 21.3 | EPI_ISL_404895 | 19-Jan-2020 | >hCoV-19/USA/WA1/2020 |
| 21 | A23403G | 28.4 | EPI_ISL_416327 | 28-Jan-2020 | >hCoV-19/Shanghai/SH0014/2020 |
| 22 | C24034T | 1.3 | EPI_ISL_410301 | 13-Jan-2020 | >hCoV-19/Nepal/61/2020 |
| 23 | G25563T | 6.2 | EPI_ISL_414625 | 26-Feb-2020 | >hCoV-19/France/PL1643/2020 |
| 24 | G25979T | 1.3 | EPI_ISL_414623 | 25-Feb-2020 | >hCoV-19/France/GE1583/2020 |
| 25 | G26144T | 8 | EPI_ISL_406036 | 22-Jan-2020 | >hCoV-19/USA/CA2/2020 |
| 26 | T26729C | 1.2 | EPI_ISL_408484 | 15-Jan-2020 | >hCoV-19/Sichuan/IVDC-SC-001/2020 |
| 27 | C27046T | 5.4 | EPI_ISL_413565 | 24-Feb-2020 | >hCoV-19/Netherlands/Berlicum_1363564/2020 |
| 28 | G28077C | 1.2 | EPI_ISL_408484 | 15-Jan-2020 | >hCoV-19/Sichuan/IVDC-SC-001/2020 |
| 29 | T28144C | 32.3 | EPI_ISL_406801 | 5-Jan-2020 | >hCoV-19/Wuhan/WH04/2020 |
| 30 | C28657T | 1.3 | EPI_ISL_414623 | 25-Feb-2020 | >hCoV-19/France/GE1583/2020 |
| 31 | T28688C | 3.1 | EPI_ISL_412981 | 18-Jan-2020 | >hCoV-19/Wuhan/HBCDC-HB-05/2020 |
| 32 | C28863T | 1.3 | EPI_ISL_414623 | 25-Feb-2020 | >hCoV-19/France/GE1583/2020 |
| 33 | G28878A | 1.2 | EPI_ISL_416317 | 25-Jan-2020 | >hCoV-19/Shanghai/SH0003/2020 |
| 34 | G28881A | 11.2 | EPI_ISL_413565 | 24-Feb-2020 | >hCoV-19/Netherlands/Berlicum_1363564/2020 |
| 35 | G28882A | 11.2 | EPI_ISL_413565 | 24-Feb-2020 | >hCoV-19/Netherlands/Berlicum_1363564/2020 |
| 36 | G28883C | 11.2 | EPI_ISL_413565 | 24-Feb-2020 | >hCoV-19/Netherlands/Berlicum_1363564/2020 |
| 37 | C29095T | 1.7 | EPI_ISL_406030 | 10-Jan-2020 | >hCoV-19/Shenzhen/HKU-SZ-002/2020 |
| 38 | G29553A | 1.4 | EPI_ISL_414616 | 8-Mar-2020 | >hCoV-19/USA/WA-UW29/2020 |
| 39 | C29635T | 1.3 | EPI_ISL_412969 | 10-Feb-2020 | >hCoV-19/Japan/Hu_DP_Kng_19-027/2020 |
| 40 | G29742T | 4 | EPI_ISL_412981 | 18-Jan-2020 | >hCoV-19/Wuhan/HBCDC-HB-05/2020 |
| 41 | G29742A | 4 | EPI_ISL_416316 | 25-Jan-2020 | >hCoV-19/Shanghai/SH0002/2020 |
| | | | EPI_ISL_416317 | | >hCoV-19/Shanghai/SH0003/2020 |

**Supplementary Table S3. Chronological order of the first appearance of mutations in SARS-CoV-2.** The 41 substitutions/mutations (including G29742A and G29742T) in the SARS-CoV-2 genomes are shown with the date of collection and geographic location of the first reported sequence.
